# RCE-IFE: Recursive Cluster Elimination with Intra-cluster Feature Elimination

**DOI:** 10.1101/2024.02.28.580487

**Authors:** Cihan Kuzudisli, Burcu Bakir-Gungor, Bahjat Qaqish, Malik Yousef

## Abstract

The computational and interpretational difficulties caused by the ever-increasing dimensionality of biological data generated by new technologies pose a major challenge. Feature selection (FS) methods aim to reduce the dimension, and feature grouping has emerged as a foundation for FS techniques that seek to detect strong correlations among features and the existence of irrelevant features. In this work, we develop Recursive Cluster Elimination with Intra-Cluster Feature Elimination (RCE-IFE), a method that iterates clustering and elimination steps in a supervised context. Recursively, feature clusters are formed, then scored, and less contributing clusters are eliminated. Next, low-scoring features in retained clusters are eliminated. Intra-cluster feature elimination aims to reduce noisy features while keeping a minimum number of predictive features. The performance of RCE-IFE is evaluated and compared to other FS techniques in several datasets. The results show that the proposed strategy effectively reduces the size of the feature set and also improves the model performance.

## 1. Introduction

Gene expression data typically come with a relatively small number of samples accompanied by a huge number of genes. This characteristic makes data processing and analysis a challenging task and is considered to be a hot topic in bioinformatics [1,2]. There is a need for a dimensionality reduction technique as a computational tool and this is where feature selection (FS) comes into play. FS refers to selection of a subset of relevant features and exclusion of redundant, irrelevant and noisy features. FS is a prerequisite for gene analysis due to the existence of redundancy and noise in gene expression data [3]. Many FS techniques are available in the literature and widely used in gene subset extraction or disease classification [4]. Recursive cluster elimination based on support vector machine (SVM-RCE) proposed by Yousef et al. [5] introduced *recursive cluster elimination* term into the literature and this approach overcame Support Vector Machines with Recursive Feature Elimination (SVM-RFE) [6], which was widely accepted as an effective approach in the field. The superiority of SVM-RCE stems from the consideration of feature (i.e., gene) clusters instead of individual features in the classification task. In SVM-RCE, samples are divided into training and test sets. Then, features are grouped into clusters. Next, feature clusters are scored according to their predictive ability using linear SVM. A predefined fraction of clusters with the lowest scores are eliminated. After each elimination step, the remaining features are fed into a classifier to quantify classification accuracy. This way, the performance of different numbers of clusters is determined as the algorithm proceeds. All the steps from clustering to cluster elimination are repeated recursively. The number of clusters formed in each iteration is a predetermined decreasing sequence.

In this study, we extend SVM-RCE and integrate elimination of features inside surviving clusters. Through this newly added step, not only less contributing clusters are dropped, but also less contributing features inside each cluster are excluded. This addition leads to deeper dimensionality reduction, thereby allowing enhancement in feature subset quality, classification performance and running time. This study proposes Recursive Cluster Elimination with Intra-Cluster Feature Elimination (RCE-IFE) that is capable of tackling large scale datasets. We compare the dimensionality reduction capability and predictive power of RCE-IFE to state-of-the-art FS methods and classifiers.

The remainder of the paper is organized as follows: Section 2 presents the main characteristics of datasets used in the study and then introduces the proposed approach in detail. In Section 3, comparative studies are described and their results presented. Section 4 concludes with possible directions for future research.

## 2. Materials and Methods

### 2.1 Datasets

We conducted our experiments on twenty datasets, consisting of gene expression, miRNA and methylation datasets accessed from GEO and TCGA databases [7,8]. Each dataset has two class labels, i.e., positive and negative, with a particular number of samples.

### 2.2 Proposed Algorithm

Our proposed algorithm, called RCE-IFE, is an iterative approach that involves removal of weakly-scoring clusters followed by intra-cluster elimination, i.e., exclusion of low-scoring features in surviving clusters. In our proposed method, the initial and desired number of clusters should be specified by the user. The input data is a two-class dataset D represented by a p x n matrix with samples represented by rows S= [s1, s2, …, sp] and features (i.e., genes) represented by columns G= [g1, g2, …, gn]. Each sample belongs to labels of pos or neg, indicating whether it is positive or negative. To begin with, the algorithm first divides the samples as training (90%) and test (10%) data and we don’t extract our test data out of the trained data, i.e., test data is kept blind so that we prevent bias in the model. Subsequently, training data undergo several steps for model construction. Once the training is completed and the model is created, testing data is input to the classifier and the classification accuracy is measured. The main steps for RCE-IE are Grouping (G), Scoring (S), Intra-cluster Feature Elimination (IFE), and Modeling (M) and its workflow is illustrated in Fig. I. Note that training data is employed through all steps and test data is essentially exploited for evaluation of model’s performance. The details of these steps are explained next.

**Fig. I.**
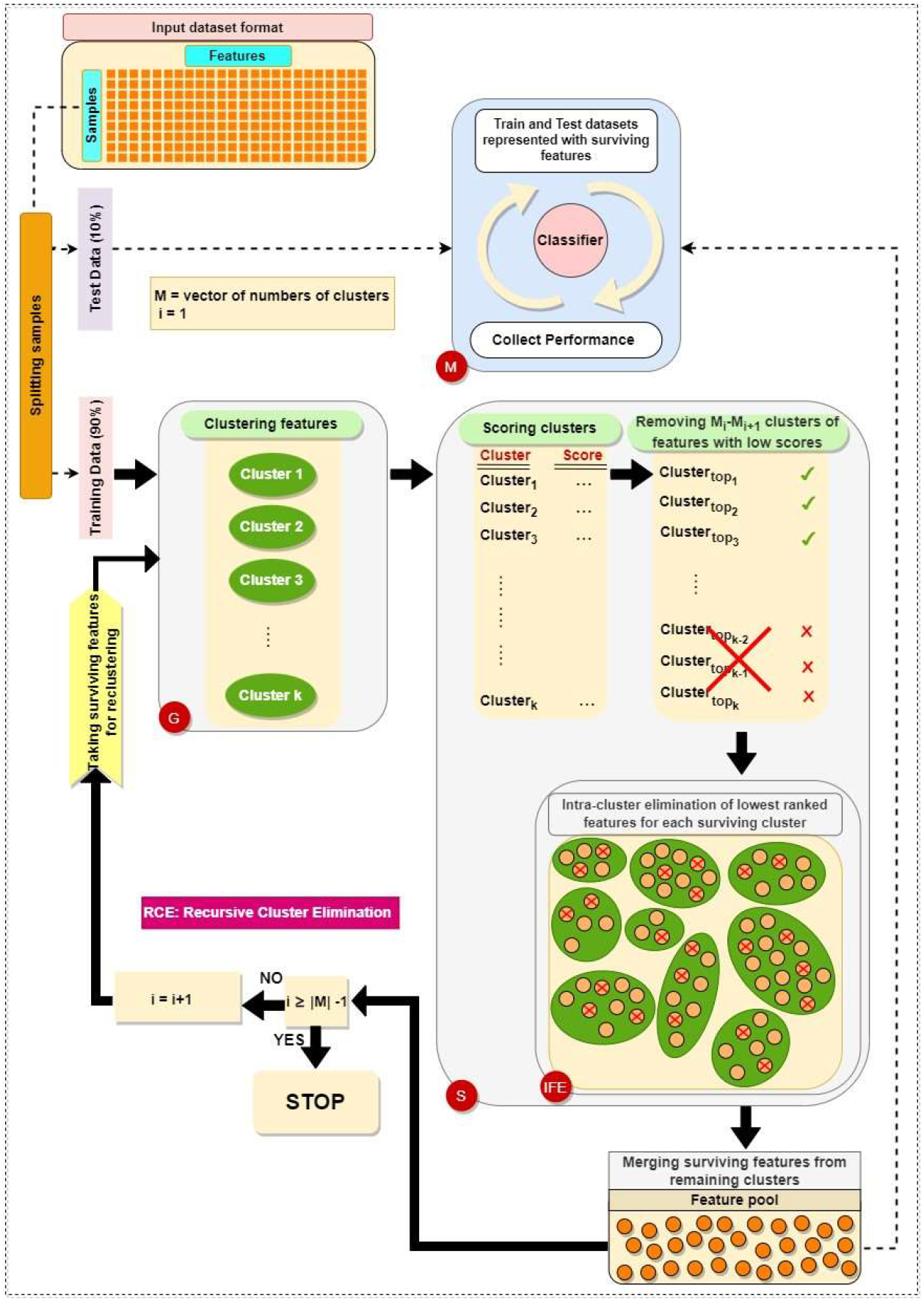
The workflow of the proposed approach

#### G step

Grouping is the initial procedure applied to training data. In this step, grouping can be done using different clustering algorithms requiring number of clusters beforehand. In our tool, K-means is applied to group genes into k clusters, which is a popular unsupervised clustering algorithm. Let C = {c_1_, c_2_, …, c_k_} represent the set of gene clusters. Each c_i_ ∈ C includes genes with highest similarity between each other and dissimilarity with genes in other clusters.

After grouping genes into clusters, we create two-class subdatasets based on gene clusters. A subdataset is actually a set of genes in a specific cluster accompanied by their respective sample values and class labels belonging to these values. Consequently, we have a subdataset for each gene cluster and k subdatasets in total. Fig. II shows the procedure for extraction of a two-class subdataset from a dataset consisting of ten samples and thirteen genes along with class labels, where pos and neg denote positive and negative class, respectively. As shown in Fig. II, the original dataset (upper left) contains genes as columns and samples as rows as mentioned before. The class column includes labels for each sample. In this scenario, there are four gene clusters (upper right) and this means we need to create four subdatasets in the end.

**Fig. II.**
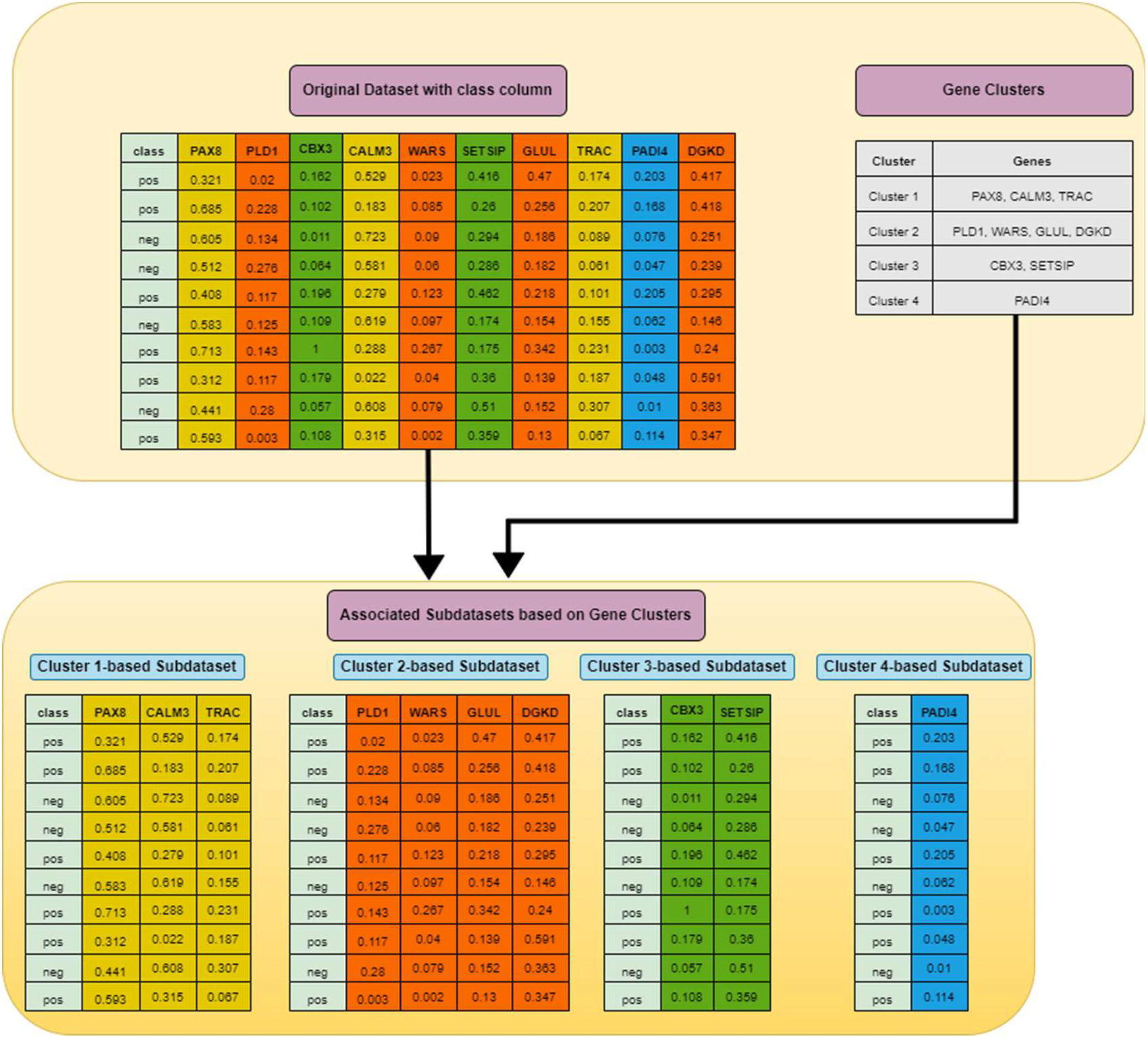
Extraction of subdatasets based on gene clusters

As illustrated in Fig. II, extraction of subdatasets is performed by selecting genes peculiar to each gene group along with their corresponding sample values and the column involving class labels. These four subdatasets are fed into S step for scoring, ranking and elimination.

#### S step

After the generation of subdatasets based on gene clusters in G step, the next step is to assign a score for each subdataset using a supervised learning algorithm such as Support Vector Machine (SVM) or Random Forest (RF). For a given subdataset, it is separated into training and test, where training data is used for learning the model and the model is validated on test data. This process is repeated t times and the classification accuracies acquired over partitions are averaged to determine the score of the subdataset. During partitioning, stratified sampling is applied rather than random sampling so that each partition has almost the same proportion of class labels as the original dataset. To put it simply, scoring a subdataset is accomplished through randomized stratified t-fold cross validation [9]. The scoring step is depicted in Fig. III.

**Fig. III.**
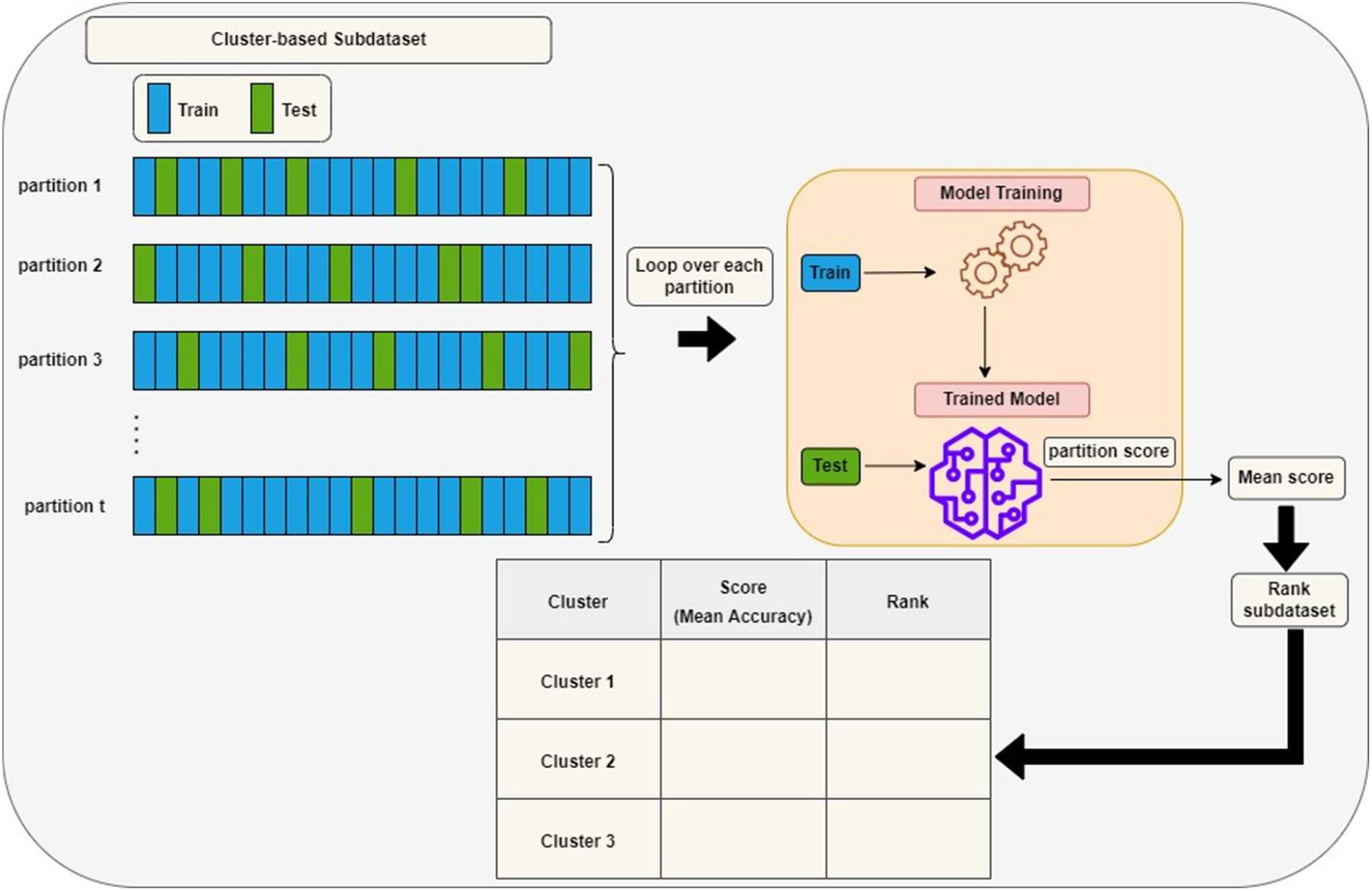
Assigning a score to a subdataset

**Fig. IV.**
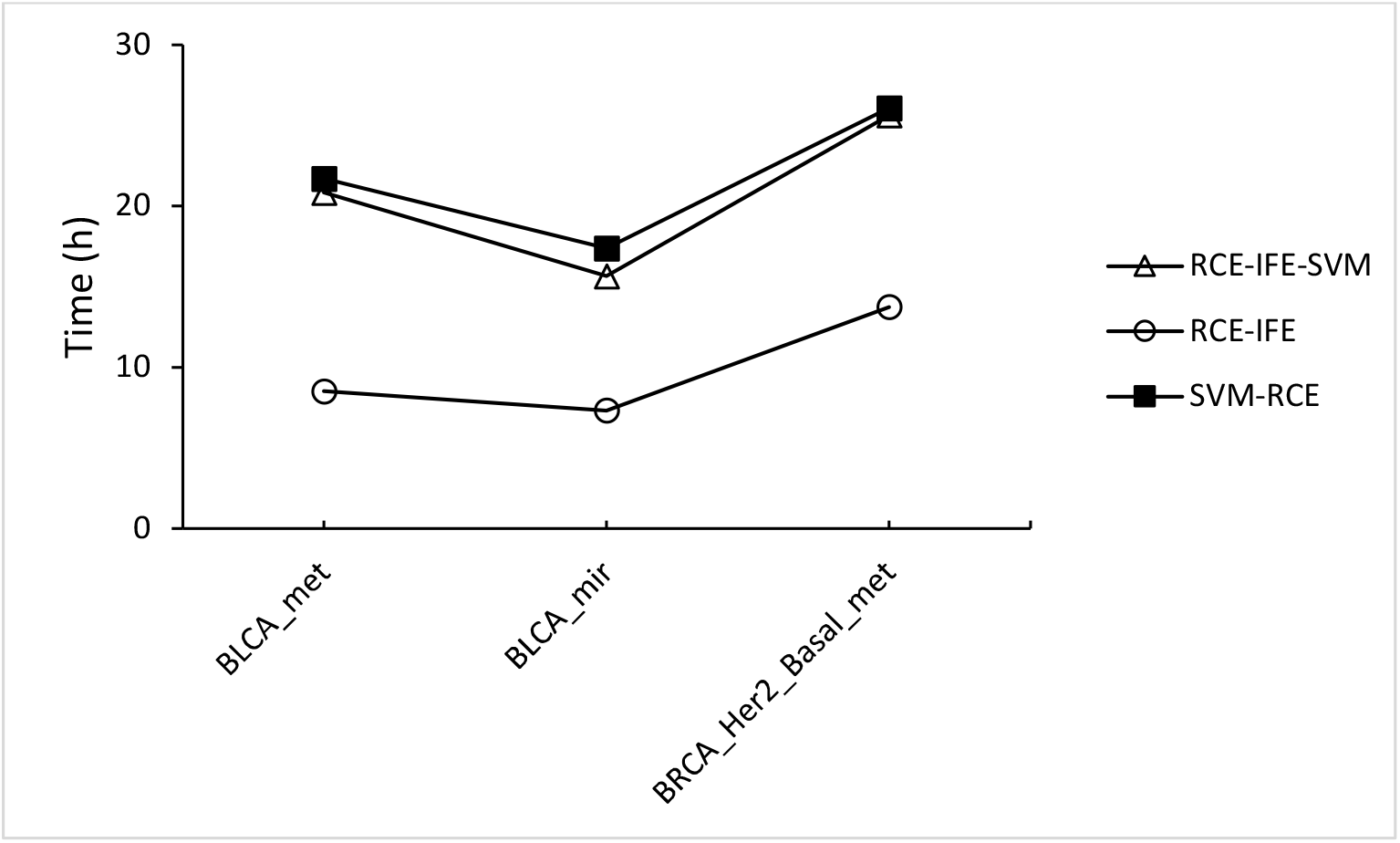
Running time comparison of RCE-IFE-SVM, RCE-IFE and SVM-RCE

Accuracy is considered among different evaluation metrics (F1 Score, Area Under the Curve (AUC), etc.) in order to score subdatasets. Once these subdatasets are assigned scores, they are ranked in descending order and d% of those with the lowest scores are eliminated.

#### IFE step

Our algorithm not only eliminates low-scoring clusters but also removes weakly-scoring genes inside surviving clusters. Scores for genes are mainly weights or coefficients determined by the preferred classification algorithm. The rate of gene removal should be ascertained in advance. IFE plays a very important role as it reduces the number of genes at each cluster elimination step. This novel process leads to further level exclusion of irrelevant / redundant genes, which is, in turn, expected to help in enhancing generalization capability of the model and obtain the same (or improve) accuracy with less number of genes. In addition, IFE will provide the most significant clusters with minimum number of significant genes in the end.

#### M step

Following the elimination of clusters with low scores and intra-cluster feature elimination of low-scoring genes in remaining clusters, all genes within remaining clusters are pooled. Genes in this pool are used to extract the training and test datasets as explained in G step. Finally, the training dataset is used to build the model and the model is tested using the test dataset. During an iteration, results (number of genes, accuracy, specificity, sensitivity, AUC, etc.) are collected at every step of cluster elimination. For instance, suppose that the algorithm starts its operation with 100 clusters initially and its elimination rate (d) is 10%. Then, our algorithm collects results for 90 clusters, 81 clusters, 72 clusters, …, and so on. With multiple iterations, results are averaged for each cluster number and evaluation of a set of clusters is achieved in this way.

The proposed method terminates when the number of clusters initially set up is less than the desired number of clusters. Initial number of clusters and desired number of clusters should be predefined input parameters like cluster and gene elimination rate.

## 3. Results and Discussion

This section presents performance of the proposed RCE-IFE approach on the conducted experiments and discusses the acquired results. We have done experiments on evaluation of RCE-IFE by comparing it with SVM-RCE on 20 publicly aforementioned datasets.

### 3.1 Experimental Setup

We have performed our analyses on Knime [10], an open-source software, due to its simplicity and support for graphical representations. In addition, Knime is an extremely integrative tool that allows integration of different scripts in many coding languages.

To ensure balance in datasets, downsampling is applied to remove the samples from the majority class while preserving all of the samples in the minority class. All methods were executed for 100 iterations to provide stability in results. We adopted a set of fixed cluster levels [90, 80, 70, 60, 50, 40, 30, 20, 10, 5, 2, 1] in the experiments. The number of genes to be removed from surviving clusters is set to be 10% if the cluster contains more than five genes. Among several performance metrics, AUC was used to evaluate classifier performances except for Section 3.4 due to the comparison with a study in the field. All experiments were conducted on a hardware environment with Intel Core i9-9900, 3.10 GHz, 64.0 GB RAM.

### 3.2 Comparison with SVM-RCE

In this section we compare our proposed approach RCE-IFE to SVM-RCE in terms of classification performance, the number of genes, and execution time. For RCE-IFE, we employed two classifiers, RF and SVM, individually. In other words, the steps in scoring clusters and genes in surviving clusters were achieved through RF and SVM separately. We use RF in RCE-IFE and denote the approach for integration of SVM into RCE-IFE as RCE-IFE-SVM. We considered the results of two clusters among various cluster numbers. Table I shows the results on the datasets. First column corresponds to AUC values followed by the number of features found by each technique. The last row refers to average AUC and feature size values obtained across all the datasets for all algorithms. We can observe that all methods achieve similar AUC values, where SVM-RCE is outperformed by RCE-IFE-SVM and RCE-IFE by 1% and 2%, respectively. However, there’s a huge difference in the number of selected features between SVM-RCE and RCE-IFE methods. RCE-IFE-SVM selects roughly half as many genes as SVM-RCE whereas RCE-IFE picks almost two and a half times fewer genes than SVM-RCE. Hence, we can say our proposed approach reduces the size of the feature subset significantly while maintaining the classifier performance.

**Table 1.** Comparison with SVM-RCE: Mean AUC and mean size of the feature subsets on 20 datasets.

|  | AUC |  |  | #Features |  |  |
| --- | --- | --- | --- | --- | --- | --- |
|  | SVM-RCE | RCE-IFE-SVM | RCE-IFE | SVM-RCE | RCE-IFE-SVM | RCE-IFE |
| GDS1962 | 0.98 ± 0.05 | 0.99 ± 0.03 | 0.98 ± 0.05 | 76.39 ± 30.23 | 33.93 ± 14.64 | 28.52 ± 10.76 |
| GDS2547 | 0.80 ± 0.11 | 0.83 ± 0.10 | 0.80 ± 0.11 | 109.05 ± 38.84 | 67.84 ± 16.57 | 51.19 ± 16.76 |
| GDS3268 | 0.80 ± 0.12 | 0.78 ± 0.12 | 0.76 ± 0.13 | 99.40 ± 40.34 | 41.44 ± 14.29 | 40.52 ± 14.45 |
| GDS3646 | 0.74 ± 0.23 | 0.77 ± 0.22 | 0.67 ± 0.25 | 38.89 ± 23.88 | 22.56 ± 9.70 | 26.44 ± 10.12 |
| GDS3837 | 0.98 ± 0.03 | 0.98 ± 0.04 | 0.98 ± 0.04 | 124.68 ± 40.48 | 62.60 ± 21.21 | 49.82 ± 16.07 |
| GDS3874 | 0.73 ± 0.20 | 0.75 ± 0.20 | 0.86 ± 0.16 | 42.19 ± 16.94 | 21.53 ± 6.32 | 24.94 ± 7.38 |
| GDS5037 | 0.73 ± 0.24 | 0.72 ± 0.22 | 0.75 ± 0.22 | 37.62 ± 14.15 | 19.00 ± 5.55 | 21.99 ± 7.39 |
| GDS5499 | 0.96 ± 0.07 | 0.95 ± 0.09 | 0.92 ± 0.10 | 179.76 ± 45.84 | 69.92 ± 14.77 | 71.34 ± 22.38 |
| GDS2519 | 0.50 ± 0.18 | 0.56 ± 0.17 | 0.55 ± 0.18 | 46.05 ± 18.44 | 32.69 ± 10.58 | 30.99 ± 9.37 |
| GDS2609 | 0.97 ± 0.14 | 0.97 ± 0.14 | 0.99 ± 0.07 | 59.03 ± 25.06 | 26.34 ± 8.96 | 19.99 ± 7.21 |
| GDS3875 | 0.84 ± 0.15 | 0.86 ± 0.14 | 0.83 ± 0.14 | 41.55 ± 14.32 | 22.22 ± 7.14 | 26.26 ± 7.95 |
| GDS3929 | 0.51 ± 0.28 | 0.48 ± 0.31 | 0.50 ± 0.26 | 17.33 ± 7.30 | 7.26 ± 2.42 | 19.82 ± 6.80 |
| GDS4228 | 0.48 ± 0.26 | 0.46 ± 0.27 | 0.80 ± 0.20 | 12.17 ± 6.18 | 6.63 ± 2.68 | 21.20 ± 6.73 |
| GDS4824 | 0.95 ± 0.22 | 0.95 ± 0.22 | 0.90 ± 0.30 | 61.57 ± 24.17 | 23.32 ± 8.39 | 14.38 ± 5.87 |
| GDS5093 | 0.90 ± 0.22 | 0.91 ± 0.22 | 0.90 ± 0.22 | 42.35 ± 23.24 | 15.40 ± 9.09 | 18.15 ± 7.56 |
| GSE15573 | 0.96 ± 0.12 | 0.94 ± 0.14 | 0.93 ± 0.17 | 63.67 ± 24.61 | 25.52 ± 8.08 | 19.25 ± 6.02 |
| GSE157103 | 0.94 ± 0.10 | 0.97 ± 0.07 | 0.93 ± 0.11 | 95.63 ± 35.05 | 42.30 ± 14.18 | 31.76 ± 11.15 |
| BLCA_met | 0.98 ± 0.06 | 0.98 ± 0.05 | 0.97 ± 0.08 | 73.81 ± 28.14 | 37.65 ± 13.28 | 27.15 ± 10.57 |
| BLCA_mir | 0.97 ± 0.08 | 0.98 ± 0.06 | 0.97 ± 0.08 | 109.19 ± 107.69 | 49.86 ± 42.42 | 25.95 ± 22.90 |
| BRCA_Her2<br>Basal met | 0.98 ± 0.05 | 0.98 ± 0.04 | 0.99 ± 0.03 | 144.27 ± 49.87 | 62.33 ± 21.95 | 53.95 ± 18.80 |
| **Average** | **0.83 ± 0.15** | **0.84 ± 0.14** | **0.85 ± 0.15** | **73.73 ± 30.74** | **34.52 ± 12.61** | **31.18 ± 11.31** |

As far as running time is concerned, RCE-IFE and RCE-IFE-SVM algorithms have less time complexity than SVM-RCE as illustrated for the given datasets in Fig IV. Note that our proposed approach might seem to have a trade-off between an additional step of internal elimination and additional gene removal. However, gene removal means dimensionality reduction and outweighs integration of internal elimination step in terms of execution time. In Fig IV, we observe that SVM-RCE is slower than RCE-IFE-SVM to a certain degree due to the contribution of internal elimination step. On the other hand, RCE-IFE is the fastest algorithm by far. It’s also noteworthy here to state that all algorithms achieve the same AUC performance (98%) on average for overall datasets with different execution times.

## Conclusion

In this paper, we address the challenge of FS task using a feature grouping-based strategy. We propose RCE-IFE that involves cluster elimination followed by removal of genes from surviving clusters to obtain non-redundant and strongly relevant features in the end. We adapted SVM to RCE-IFE and represented it as RCE-IFE-SVM. For comparison purpose, we employed RCE-IFE, RCE-IFE-SVM and SVM-RCE. Experiments show that although RCE-IFE and RCE-IFE-SVM methods exhibit slightly better classification performance than the crude method, SVM-RCE, they accomplish it with a higher dimensionality reduction rate, i.e., with significantly less number of features. Hence, IFE proves to be an effective step in dimensionality reduction without deteriorating classifier performance. Furthermore, RCE-IFE and RCE-IFE-SVM approaches are faster than SVM-RCE, which implies another contribution of IFE. However, RCE-IFE is much faster than RCE-IFE-SVM. Overall, results show that RCE-IFE outperforms SVM-RCE in terms of number of features, performance and running time.

